# The role of the *fruitless* P3 transcript in development in *Drosophila melanogaster*

**DOI:** 10.64898/2026.11.03.764014

**Authors:** Sara A. Kulinski, William Yeung, Amanda J. Moehring

## Abstract

The generation of multiple transcripts from a single gene through alternative promoter usage and alternative splicing is a major source of functional diversity in eukaryotic genomes. The *fruitless* (*fru*) gene of *Drosophila melanogaster* is a classic example of transcript complexity, producing at least 15 transcript isoforms through the use of alternative promoters and terminal exons. While transcripts derived from the sex-specifically spliced P1 promoter have been extensively studied for their roles in sex determination and courtship behaviour, the functions of most non-sex-specific *fru* transcripts remain poorly understood. Previous studies suggested that transcripts originating from the P3 and/or P4 promoters may contribute to developmental viability, but their individual roles could not be resolved because available mutations disrupted multiple transcript classes simultaneously. Here, we used transcript-specific RNA interference (RNAi) and the Gal4/UAS system to investigate the function of *fru* P3 transcripts. Loss of P3 expression significantly increased developmental time to pupariation and resulted in complete lethality during pupal development despite normal survival to pupariation. Most mortality occurred during early and mid-pupal stages, indicating a critical requirement for P3-derived transcripts during metamorphosis. Notably, P3 knockdown did not phenocopy complete loss of *fruitless*, as knockdown individuals survived substantially later into development than *fru* null mutants, suggesting that additional *fruitless* transcript classes contribute essential functions during earlier developmental stages. Together, these findings establish an independent developmental role for the *fru* P3 transcript class and demonstrate that transcript-specific functions contribute substantially to the biological complexity of the *fruitless* locus.

## INTRODUCTION

Alternative splicing and alternative promoter usage are major sources of regulatory and functional diversity in eukaryotic genomes, allowing a single gene to generate multiple transcript and protein isoforms with distinct developmental, physiological, and behavioural functions (Breitbart et al. 1987; Chen and Manley 2009). Through the production of alternative transcripts, genes can expand their functional repertoire, increasing regulatory and phenotypic diversity without a corresponding increase in gene number. Genes with extensive alternative promoter usage and alternative splicing are thought to have acquired their transcript diversity through the accumulation of novel regulatory and coding elements over evolutionary time (Ayoubi and Van de Ven 1996; Keren et al. 2010). Yet reconstructing this evolutionary process requires knowledge of the functions associated with each transcript class. In the absence of functional information, it is difficult to determine whether newly evolved transcripts acquired novel roles, partitioned ancestral functions, or modified pre-existing developmental pathways. Understanding the functions of individual transcript classes is therefore essential for explaining not only the biology of complex genes, but also how such complexity evolves.

In *Drosophila melanogaster*, genes involved in neural development and behavior are particularly enriched for alternative transcript usage. For example, *paralytic*, which encodes a voltage-gated sodium channel required for neuronal action potentials, generates dozens of transcript isoforms through extensive alternative splicing (Thackeray and Ganetzky 1994; Park and Graveley 2007; Öztürk-Çolak et al. 2024), while the protein kinase gene *foraging* produces multiple transcripts that contribute to variation in feeding and locomotor behaviours (reviewed in Anreiter and Sokolowski 2019). These heavily alternatively spliced genes provide valuable models for understanding how transcript diversity contributes to complex phenotypes. One of the best-studied alternatively spliced genes is the transcription factor gene *fruitless* (*fru*) (Ito et al. 1996; Ryner et al. 1996). The *fru* locus produces at least 15 alternative transcripts through a combination of one of five promoter-associated alternative first exons (P1-P5) and four alternative terminal exons (A-D), with a series of shared central common exons (Ito et al. 1996; Ryner et al. 1996). These transcripts encode proteins that contain an N-terminal BTB domain and C-terminal zinc-finger DNA-binding motifs characteristic of transcriptional regulators (Ito et al. 1996; Ryner et al. 1996). Historically, studies of *fru* have focused almost exclusively on transcripts derived from the P1 promoter due to their sex-specific alternative splicing and role in male sex determination and courtship behaviour (Ryner et al. 1996; Heinrichs et al. 1998; Demir and Dickson 2005; Yamamoto and Koganezawa 2014), and far less is known about the other transcripts produced from *fru*. Recently, transcripts derived from the P2 promoter were found to affect female receptivity, demonstrating that some of the non-sex-specifically spliced transcript classes also have a role in sexual development and behaviour (Chowdhury et al. 2020).

Mutations affecting combinations of transcript classes, including those simultaneously disrupting P1, P2, and P3 transcripts or disrupting both P3 and P4 transcripts, result in developmental defects and lethality during metamorphosis (Anand et al. 2001). In contrast, flies lacking only P1- or P2-derived transcripts remain viable, suggesting that the developmental defects observed in these broader mutants likely arise from disruption of P3 and/or P4 transcript classes (Anand et al. 2001). However, because existing mutations affect multiple promoters simultaneously, it remains unclear whether either transcript class independently contributes to viability or whether lethality results from the combined disruption of several transcript classes. Expression patterns suggest that P3 and P4 isoforms may participate in developmental processes necessary for successful metamorphosis and adult emergence, although direct functional evidence has been lacking (Leader et al. 2018). Because the *fruitless* P3 and P4 promoters are located closer to the central shared coding region of the locus than the more distal upstream promoters, and appear to play a role in critical developmental processes, they may represent transcript classes associated with older components of the gene’s evolutionary history. If so, their functions could provide insight into the ancestral roles of *fruitless* prior to the evolution of additional promoter-specific transcripts involved in specialized reproductive and behavioral processes. Characterizing the developmental functions of these poorly understood transcript classes therefore has implications beyond understanding *fruitless* itself, offering a framework for investigating how alternatively spliced genes originate, diversify, and evolve increasingly complex regulatory architectures over evolutionary time.

One major obstacle has been the complex structure of the *fruitless* locus. Because multiple transcript classes share coding regions and regulatory elements, mutations designed to disrupt one promoter often affect additional transcripts. As a result, it has been difficult to determine the independent contributions of individual transcript classes to development and viability. The Gal4/UAS system coupled with transcript-specific RNA interference (RNAi) provides a powerful means of overcoming the genetic complexity of the *fruitless* locus (Brand and Perrimon 1993). By selectively reducing expression of a single transcript class while leaving the remainder of the gene intact, we can evaluate the function of individual transcripts.

In this study, we used transcript-specific RNAi to investigate the developmental functions of the *fruitless* P3 transcripts. P3 knockdown delayed larval development, significantly increased time to pupation, and resulted in complete lethality during the pupal stage despite normal survival to pupariation. Notably, knockdown of P3 transcripts did not recapitulate the phenotype of *fru* null mutants, as P3 knockdown individuals survived to the pupal stage, whereas flies lacking the entire *fruitless* locus died earlier in development prior to pupariation. Together, these findings demonstrate that P3 transcripts have an essential and previously unresolved role in developmental viability independent of other *fruitless* transcript classes and reveal an important developmental function for non-sex-specific *fruitless* isoforms.

## METHODS

### Fly maintenance and stocks

All transgenic *D. melanogaster* were acquired from the Bloomington *Drosophila* Stock Center (BDSC; Bloomington, Indiana) or made within our laboratory. The BDSC stocks are as follows: *Act5C-*Gal4 (stock #3954: *y^1^ w*; P{w^+mC^=Act5c-Gal4}17bFO1/TM6B,Tb^1^*), balancer (#3703: *w^1118^/Dp(1;Y)y^+^; CyO/nub^1^ b^1^ sna^Sco^ lt^1^ stw^3^; MKRS/TM6B,Tb^1^*), phiC31 integrase (#25709: *y^1^ v^1^ P{y^+t7.7^=nanos-phiC31\int.NLS}X; P{y^+t7.7^=CaryP}Msp300^attP40^*), and *fru* knockout (#66692: *Df(3R)fru^4-40^, fru^4-40^/TM6B,Tb^1^*). The wildtype strain, BJS, was obtained from the Sinclair Laboratory, Western University. All strains, injected embryos, and assays (see below) were conducted on standard cornmeal-agar media (BDSC recipe) at 25°C, ∼70% relative humidity, and a 14h:10h light:dark cycle.

### shRNA design

We originally aimed to create RNAi knockdowns for both P3 and P4 transcripts. To design appropriate targeting constructs, we first sequenced the P3 and P4 first exon regions in the fly stock used to create the RNAi lines (#25709), the balancer stock (#3703) and the *Act5c*-Gal4 stock (#3954), using the following primers: P3-F: 5’CACATGCGAAATTCACCGCT, P3-R: 5’ACCGACATTCCTTTCTGTTTCT, P4-F: 5’GGACAGTGCTGCGTTCCTTA, P4-R: 5’CTCCGGTGGGTCTTGTGATG. DNA was purified using the Monarch DNA Gel Extraction Kit Protocol (NEB #T1020) and sequenced at Robarts Research Institute (London, ON).

The targeting sequences of P3 and P4 shRNAs were designed using InvivoGen siRNA Wizard Software. The P3 or P4 unique first exon regions of the transcripts were targeted to allow for RNAi knockdown of these transcripts without affecting other transcripts from the *fru* gene, lowering the chances of off-target effects. BLASTn was used to assess possible off-target binding. Any of the potential targeting sequences that shared 18 or more nucleotides with other transcripts in the *D. melanogaster* exome were not used. P3 and P4 “Sham” targeting sequences were also designed to use as a control: a scrambled version of their corresponding targeting sequence with no genomic or transcriptomic targets, using the same InvivoGen software and assessed for off-targeted effects as above. Hairpins were created by adding “tagttatattcaagcata” to the 5’ end of the antisense strand, then adding the complement of the antisense sequence to the 5’ end. To integrate the hairpin constructs into the selected vector, NheI and EcoRI sites were added to the 5’ and 3’ ends of the hairpins: 5’CTAGCAGT and 3’GCG added to the ends of the “top” strands, and 5’AATTCGC and 3’ACTG added to the ends of the “bottom” strand.

The P3, P4, P3-Sham, and P4-Sham shRNAs were integrated into the plasmid pVALIUM20, a gift from Dr. Norbert Perrimon and Dr. Jian-Quan Ni, following the protocol described by Chang et al. (2014). The pVALIUM20 vector contains a visible selectable marker (*vermilion* eye colour mutation rescue) for selection of successful transformants, an attB site for targeted integration of the vector using phiC31 integrase, UAS sequences, and an AmpR element (Groth et al., 2004; Isaacson et al., 2022). The vector was double digested using New England Biolabs (NEB) restriction enzymes NheI and EcoRI. NEB Ligation protocol with T4 DNA ligase and high efficiency transformation protocols, #M0202 and #C3019I respectively, were used to integrate the shRNA sequences into pVALIUM20 plasmids and transform the recombinant plasmid into 10-Beta *E. coli* cells following standard protocol. One hundred μL of the cell-media mixture was plated onto lysogeny broth (LB) +ampicillin plates (100µL/mL) and grown for 16 hours at 37°C. Six colonies were then selected and grown in 4 mL of LB+ampicillin (100µL/mL) for 16 hours. Plasmid DNA was extracted using the Purelink Quick Plasmid Miniprep Kit (Invitrogen). Primers flanking the shRNA insertion region (F: 5’ACCAGCAACCAAGTAAATCAAC and R: 5’TAATCGTGTGTGATGCCTACC primers) and a touchdown PCR protocol were used to PCR amplify then visualize successful shRNA insertion on an agarose gel. Successful ligation and transformation were visualized through a 350 bp RCR product, whereas unsuccessful transformants were observed as 900 bp PCR product. Successful transformants were sequenced for verification using the forward primer.

Microinjection of plasmids into embryos were performed in-house as in Yeung (2021) into the phiC31 integrase stock containing the *vermilion* eye mutation. Transformants with *vermilion* rescue were first crossed to the balancer stock then confirmed via PCR. A series of crosses were performed to create the final genotype *v-*/+; *shRNA*/*CyO*; +/+ for UAS-P3-RNAi, UAS-P3-Sham, UAS-P4-RNAi, and UAS-P4-Sham stocks. Note that homozygous shRNA individuals were viable and segregating in the stock.

### qPCR

RNA extractions were performed on samples of *D. melanogaster* at the early pupal stage, collected 24-36 hours after pupariation, following the protocol in Allen (2016). Three biological replicates (N=12 pupae each) from each of the four transgenic genotypes (UAS-P3-RNAi, UAS- P3-Sham, UAS-P4-RNAi, UAS-P4-Sham), the *Act5c-*Gal4 alone genotype, and the four genotypes with the UAS and Gal4 combined (UAS;Gal4) were extracted, for a total of 27 samples. Samples were incubated at 25°C for one hour post-collection and then flash frozen by liquid nitrogen. RNA quality and quantity were evaluated using a NanoPhotometer P300 (Implen) spectrophotometer. RNA samples with a concentration greater than 1500 ng/μL, a 260/280 ratio of ∼2.0, and 260/230 ratio above 1.8 were kept for future use. Complimentary DNA (cDNA) was synthesized using a Thermo Scientific Maxima H Minus cDNA Synthesis Kit with dsDNase (Thermofisher) in 40 μL total volume using 2 µg of RNA following the manufacturer’s protocol. cDNA was visualized using primers for mRNA of the RpL housekeeping gene (Forward: 5’GGCATCAGATACTGTCCCTTG, Reverse: 5’CCAGTCGGATCGATATGCTAA primers), to qualitatively observe consistency between samples.

Primers used for quantification mRNA expression for P3, P4, and the control gene *alpha- Tubulin* in *D. melanogaster* were designed using Primer-BLAST: *alpha-Tubulin*-F: GCGTCGGTCAATTCAATCTT, *alpha-Tubulin*-R: AAGCTGCAACCTCTTCGTCA, P3-F: TCATCAGCAAATGCCTCGTCG, P3-R: CTCCTTGGTCAGTGTTGTACCTTG, P4-F: TGGCTGCTGGAATCACTCTT, and P4-R: GGTCTGGTGAGCCTTGACTG. Optimization of P3 and P4 primers was performed with a PCR thermal gradient (55–65°C). Both P3 and P4 primers didn’t have visible off-target binding or primer dimers and had an optimal annealing temperature of 58-60°C. The efficiency and specificity of both primers were confirmed with a standard curve. Wildtype BJS female *D. melanogaster* cDNA was diluted in a series of 10-fold dilutions and used in PCR reactions with P3 or P4 primers to build 5-log standard curves for each primer set. Each sample was loaded in triplicate, a no-template control added for each primer set. The PCR reaction ran for 40 cycles with an annealing temperature of 60°C using the CFX96™ Touch Real-Time PCR detection system. Efficiency, R^2^ values and Ct values for melt curves were calculated using BioRad CFX Manager Software and evaluated prior to running experimental qPCRs.

Quantitative PCR was performed to assess the expression levels of P3, P4, and the control gene *alpha-Tubulin*. Each of the three biological replicates was repeated in two technical replicates. Primer pairs were not multiplexed; amplifications were done separately but in parallel using Thermo Fisher Scientific 10x PowerUp SYBR Green Master Mix and an annealing temperature of 60°C for 39 cycles, and then one cycle with an annealing temperature of 65°C, on a CFX96™ Touch Real-Time PCR detection system. Relative and normalized expression were assessed through BIO RAD CFX Manager 3.0 software (Version 3.0.1215.0601).

### Developmental assay

The time to transition between developmental stages and number of survivors to reach each stage of development were assayed for P3-RNAi. P4 RNAi was not assayed as it did not generate successful knockdown of the P4 transcript (see Results). There were four groups tested in this assay: the RNAi knockdown UAS-P3-RNAi/*CyO*;*Act5c*-Gal4/+, and three control groups of UAS-P3-Sham/*CyO*;*Act5c*-Gal4/+ sham knockdown, the UAS-P3-RNAi/*CyO* alone, and *Act5c*- Gal4/*TM6B,Tb^1^*alone.

For the UAS-P3-RNAi and *Act5c*-Gal4 control groups, 200+ flies were taken from each line and placed into their own separate embryo collection cage. To make the UAS-P3- RNAi/+;*Act5c*-Gal4/+ and UAS-P3-Sham/+;*Act5c*-Gal4/+ flies, 200 female virgin flies were collected between the age of 1-6h from the *Act5c*-Gal4/ *TM6B,Tb^1^*line and put into an embryo collection cage with either 100 homozygous males from the UAS-P3-RNAi or UAS-P3-Sham lines. After three days of acclimation in the embryo collection cage, an apple juice agar petri dish with active yeast paste was switched into the embryo collection cages and replaced every 30 minutes for two hours. After the 2-hour period, fresh plates were added and all groups were allowed to lay eggs for three hours. Between 200-250 eggs were collected from each plate and placed in groups of 40 on 22x22 mm glass cover slips. The cover slips were placed onto fresh food plates, separated by treatment group. Plated eggs were incubated for 24 hours. The total number of eggs that hatched and reached the larval state was recorded for each treatment group. 288 test tubes (12x75mm) were filled with standard cornmeal food (2 cm from base) 24 hours prior to loading and sorted into 4 tube racks. A total of 72 larvae were randomly chosen from each group and placed individually in a test tube filled with food. The test tube was marked with the genotype of the larva within, and using a random number generator, was given a randomized spot within the tube rack. After filling all 288 test tubes with larvae, the tube racks were placed in a 25°C incubator for the remainder of the assay.

The stage of development (larvae, pupa, or adult) was recorded for each individual every 12 hours (11am/11pm) until all larvae reached the adult stage or died. Mean time to reach each stage of development (larvae to pupae, pupae to adult), sex, and number of individuals that survived each developmental stage was recoded. As expected, approximately 50% of the progeny from the crossed lines UAS-P3-RNAi or UAS-P3-Sham to *Act5c*-Gal4/*TM6B, Tb^1^* did not possess the *Act5c*-Gal4 but the balancer chromosome containing the *Tb* visible phenotype instead and were eliminated from the assay as soon as the *Tb* phenotype was visible (late larval stage). Since each individual was stored in its own assay chamber, all of the previous data from *Tb* individuals that survived to late larval stage was able to be removed from the data set. This assay was repeated three times per treatment group, leading to 216 larvae per genotype being tested, and 864 individuals tested overall.

Once development arrested, the stage and phenotype at death for of the experimental group was recorded in detail: axis that developed (anterior versus posterior), presence of pigmentation and basic anatomical features. A *fruitless* complete knockout line (*fru^4-40^*) was scored in the same manner for comparison to P3 knockdown flies.

### Data analysis

For qPCR, expression was normalized to the reference gene (*alpha-Tubulin*) and ΔΔCt values were calculated using BioRad CFX Manager 3.0 software. Values were compared using a one-way ANOVA followed by *post hoc* t-test pairwise comparisons using a threshold of α = 0.05. A Holm-Bonferroni analysis was conducted to correct for multiple comparisons (Holm, 1979). Developmental times were evaluated using a Wilcoxon rank sum test that was conducted between groups and within each developmental group (“Time to pupation” and “Time to adult” stages). For the survival analysis, assays were scores categorically for survival (“dead” or “alive”). The proportion survived was calculated for experimental and control groups and analyzed using a Fisher’s exact test (2x2 contingency table) between each pair of genotypes. *P*- values were corrected for multiple tests using Holm-Bonferroni analysis (Holm, 1979). All statistical analyses and graphical representation of development and survival data was performed using R 4.3.1 interfaced in RStudio 2023.06.0+421.

## RESULTS

### Development time is greater in *fru* P3 knockdown strains

Expression of P3 and P4 transcripts was quantified by qPCR. Relative to the P3 RNAi genotype, P3 transcript abundance was significantly higher in all control genotypes (3.75-7.39 fold; corrected P ≤ 0.05). However, there was no significant differences in P4 transcript abundance among the P4 RNAi and control genotypes (Table 1), and thus no further tests were conducted for P4 RNAi.

**Table 1.** Summarized data of P3 or P4 transcript normalized expression from qPCR for both P3 and P4 knockdown groups and relevant controls

| **Group** | **Genotype** | **Normalized expression** | **Normalized Relative expression<sup>1</sup>** | ***P*-value<sup>1</sup>** | **Corrected *P*-value<sup>1</sup>** |
|---|---|---|---|---|---|
| P3 knockdown | UAS-P3-RNAi; *Act5c*-Gal4 | 0.00117 | 1.00000 | n/a | n/a |
|  | UAS-P3-RNAi | 0.00862 | 7.39252 | < 0.001 | < 0.001 |
|  | UAS-P3-Sham; *Act5c*-Gal4 | 0.00437 | 3.75133 | 0.0336 | 0.0336 |
|  | UAS-P3-Sham | 0.00733 | 6.28431 | 0.0063 | 0.0125 |
|  | *Act5c*-Gal4 | 0.00810 | 6.95184 | 0.0045 | 0.0136 |
| P4 knockdown | UAS-P4-RNAi; *Act5c*-Gal4 | 0.00214 | 1.00000 | n/a | n/a |
|  | UAS-P4-RNAi | 0.00241 | 1.12643 | 0.7962 | 1.2311 |
|  | UAS-P4-Sham; *Act5c*-Gal4 | 0.00236 | 1.10465 | 0.6155 | 1.2311 |
|  | UAS-P4-Sham | 0.00183 | 0.85750 | 0.3830 | 1.1489 |
|  | *Act5c*-Gal4 | 0.00408 | 1.90780 | 0.1870 | 0.7479 |

<sup>1</sup> Relative expression, raw *P*-values, and corrected (Holm-Bonferroni) *P*-values from expression data where expression of either *fru* P3 or P4 was compared between the *fru* P3 or P4 RNAi knockdown groups and controls.

Developmental time was scored and compared between *fru*-P3 knockdown (UAS-P3- RNAi; *Act5c-*Gal4) and three controls (UAS-P3-Sham;*Act5c-*Gal4, UAS-P3-RNAi, UAS-P3- Sham, and *Act5c-*Gal4) from the larval to pupal stage and the pupal to adult stage (Figure 1A, 1B; Table S1, S2). The knockdown genotype exhibited the longest larval-to-pupal developmental time (median = 144 h), significantly exceeding all control genotypes (120-132 h; corrected *P* < 0.001). No *fru* P3 knockdown pupae reached the adult stage, preventing comparisons of developmental time between groups for pupal to adult stage.

**Figure 1.**
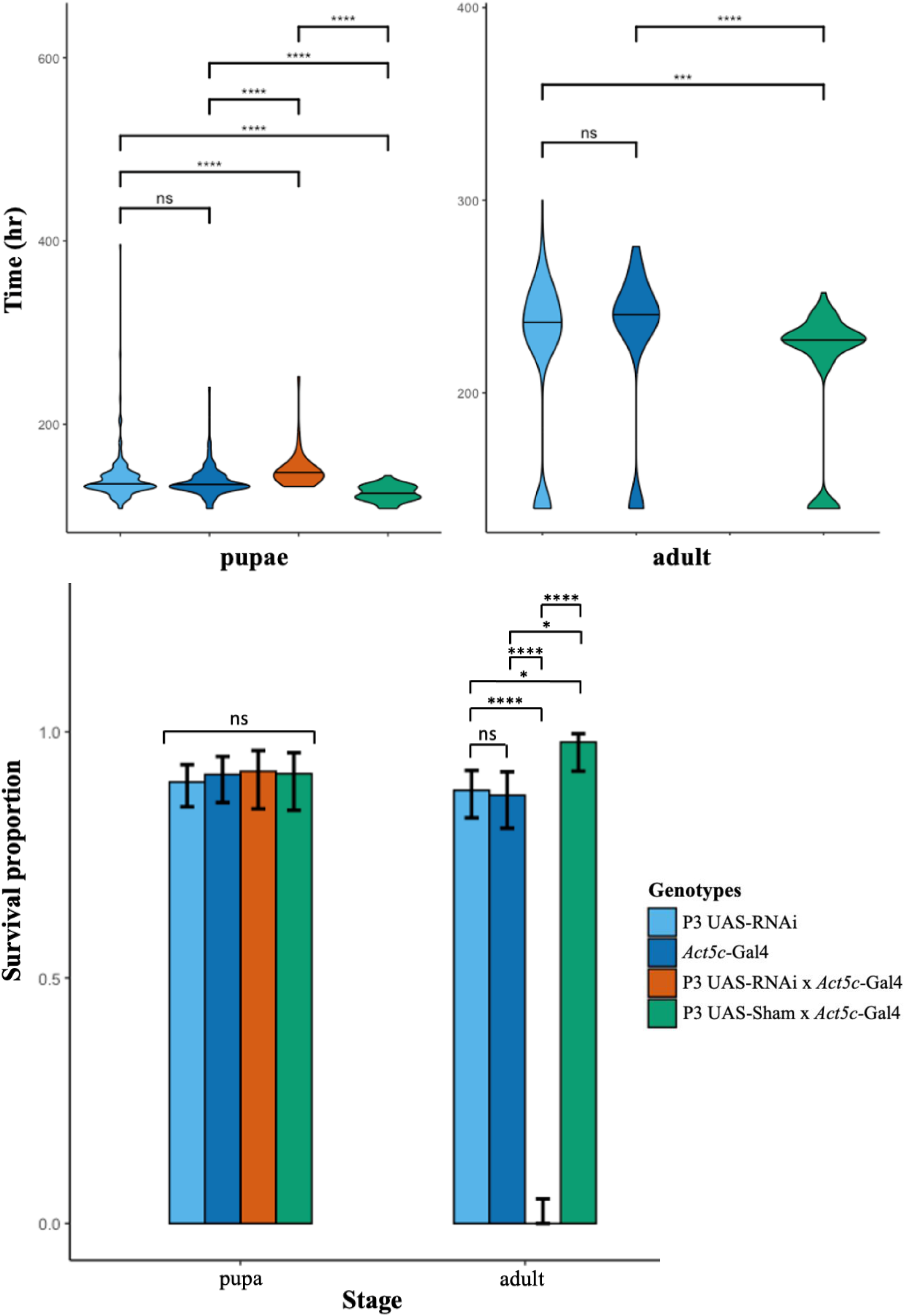
Developmental time and survival are impacted by loss of *fru* P3. **(A)** Time (hours) from egg hatching to the beginning of pupation, **(B)** time (hours) from the beginning of pupation to eclosion as adults, and **(C)** proportion surviving to pupariation and adults for P3 knockdown (UAS-P3-RNAi;*Act5c-*Gal4) in orange and three controls of UAS-P3- RNAi (light blue), UAS-P3-Sham;*Act5c-*Gal4 (green), and *Act5c-*Gal4 (dark blue). Note that no UAS-P3-RNAi;*Act5c-*Gal4 flies survived to adults. The median of each group is indicated by the horizontal line through each of the sample plots in A and B. Error bars represent a 95% confidence interval in C. *P* values indicated as *\* P*<0.05, ** *P*<0.01, *** *P*<0.001, and **** *P*<0.0001.

### Knockdown of *fru* P3 results in lethality during the pupal stage

Survival from larva to pupa did not differ significantly among genotypes, with 89.8- 92.0% of individuals reaching pupariation (*P* > 0.05; Figure 1C; Table S2, S3). In contrast, no P3 knockdown individuals survived to adulthood, whereas 79.2-89.6% of control animals successfully eclosed (corrected *P* < 0.0001).

### Knockdown of *fru* P3 causes lethality later than complete loss of *fruitless*

To determine when lethality occurred, dead P3 knockdown individuals were categorized according to pupal developmental stage (Table S4). Most deaths occurred during early to mid- pupal development, with 43.7% of individuals dying during the stage characterized by developing structures but no visible pigmentation and 33.3% dying during the prepupal stage (1- 12 h after pupariation). Fewer individuals died during pigment development (17.3%) or later pupal stages after major adult structures had formed (5.8%). In contrast, no individuals carrying the *fruitless* null allele (*fru^4-40^*) survived to pupariation, indicating that knockdown of P3 transcripts results in lethality substantially later in development than complete loss of *fruitless*.

## DISCUSSION

Alternative promoter usage and alternative splicing can generate substantial functional diversity within a single gene, but understanding how that diversity contributes to organismal biology requires characterization of individual transcript classes (Breitbart et al. 1987; Chen and Manley 2009; Keren et al. 2010). Although *fruitless* (*fru*) has long served as a model for studying the genetic control of behavior, most investigations have focused on the sex-specifically spliced P1 transcripts that regulate male courtship and reproductive behaviors (Ryner et al. 1996; Demir and Dickson 2005; Yamamoto and Koganezawa 2014). Consequently, the functions of many non-P1 transcript classes remain poorly understood. Here, we used transcript-specific RNA interference (RNAi) to investigate the role of the *fru* P3 transcript class and found that reduction of P3 expression resulted in delayed development and complete pupal lethality. These findings demonstrate that P3-derived transcripts perform an essential developmental function. Furthermore, they demonstrate that another transcript of *fru*, likely *fru*-P4, plays a critical role earlier in development, prior to pupariation.

Expression data provide additional support for a developmental role for P3 transcripts. During larval development, P3 transcripts are most highly expressed in the central nervous system and trachea (Leader et al. 2018), two organ systems that undergo extensive remodeling during pupal metamorphosis when larval tissues are eliminated or reorganized while adult structures differentiate and mature (Ashburner 2005). Given the established role of Fru proteins as transcriptional regulators (Ito et al. 1996; Ryner et al. 1996), reduced P3 expression may alter downstream gene networks required for successful developmental progression through the extensive neural reorganization that occurs during metamorphosis. Alternatively, the high expression of P3 transcripts in the tracheal system raises the possibility that defects in respiratory system remodeling may also contribute to lethality. Consequently, the developmental defects observed following P3 knockdown may reflect disruption of one or more essential transcriptional programs in tissues where P3 expression is normally most abundant.

The present findings may also provide insight into the evolution of transcript complexity. Transcript diversity generated through alternative promoter usage and alternative splicing is thought to accumulate over evolutionary time through the acquisition of novel regulatory and coding features (Ayoubi and Van de Ven 1996; Keren et al. 2010). In such systems, different transcript classes may become specialized for distinct biological functions while remaining linked within a common genomic locus. The observation that P3 transcripts are required for developmental viability, whereas P1 and P2 transcripts primarily regulate reproductive behavior, supports the hypothesis that functional partitioning has occurred within *fruitless*. Understanding the functions of additional transcript classes may therefore help reveal how increasingly complex regulatory architectures evolve while preserving essential developmental functions.

One limitation of the present study is that the P4 RNAi construct did not significantly reduce P4 transcript abundance, preventing direct assessment of P4 function. Consequently, whether P4 transcripts contribute to developmental viability remains unresolved. An additional limitation is that P3 knockdown was achieved using a ubiquitously expressed *Act5c*-Gal4 driver, resulting in transcript reduction throughout the organism and across all developmental stages. While this approach was effective for establishing an essential role for P3 transcripts in viability, and the timepoint lethality occurs, it does not identify the tissue or developmental stage in which P3 function is required. Future studies employing tissue-specific and temporally restricted GAL4 drivers could be used to determine whether the observed lethality primarily results from loss of P3 function in the nervous system, trachea, or other tissues, and to identify the developmental window during which P3 activity is required. Such experiments would provide important insight into the mechanisms through which this transcript class contributes to metamorphic development.

In conclusion, our results demonstrate that P3-derived *fruitless* transcripts are independently required for normal development and survival through metamorphosis. Reduction of P3 expression delayed larval development and resulted in complete pupal lethality despite successful pupariation. Loss of *fru*-P3 did not recapitulate the timepoint of lethality observed with full loss of *fru*, indicating that earlier lethality results form a different subset of *fru* transcripts. These findings establish a previously unresolved developmental function for a non- sex-specific *fruitless* transcript class and provide further evidence that the biological roles of the *fruitless* locus extend well beyond the regulation of reproductive behavior.

## SUPPLEMENTARY MATERIAL

## SUPPLEMENTARY MATERIAL

**Table S1.**
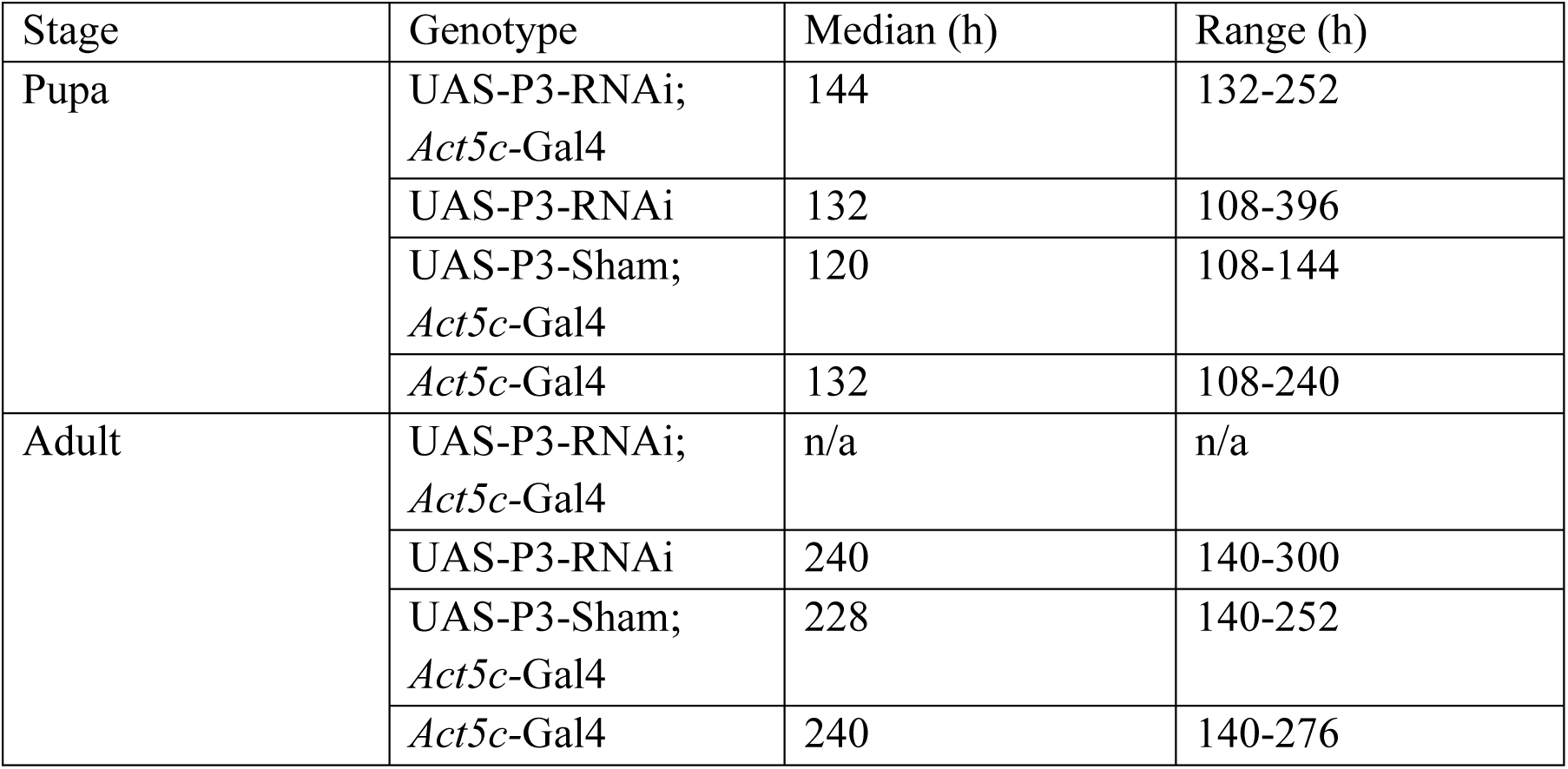
Summarized data for developmental assay showing median time to move from larval to pupal (pupa) and pupal to adult (adult) stage for each group and the range of times within each group, in hours.

**Table S2.** Raw and corrected (Holm-Bonferroni) *P*-values from developmental assays where developmental time and survival were scored. Groups separated by assay type (developmental time vs. survival) and by stage (pupa vs. adult).

| Analysis | Stage | Comparison | *P*-value | Corrected *P*-value |
|---|---|---|---|---|
| Developmental Time – Wilcoxon | Pupa | (UAS-P3-RNAi; *Act5c*-Gal4) to (UAS-P3-Sham; *Act5c*-Gal4) | < 0.0001 | < 0.0001 |
|  |  | (UAS-P3-RNAi; *Act5c*-Gal4) to (UAS-P3-RNAi) | < 0.0001 | < 0.0001 |
|  |  | (UAS-P3-RNAi; *Act5c*-Gal4) to (*Act5c*-Gal4) | < 0.0001 | < 0.0001 |
|  |  | (UAS-P3-Sham; *Act5c*-Gal4) to (UAS-P3-RNAi) | < 0.0001 | < 0.0001 |
|  |  | (UAS-P3-Sham; *Act5c*-Gal4) to (*Act5c*-Gal4) | < 0.0001 | < 0.0001 |
|  |  | (UAS-P3-RNAi) to (*Act5c*-Gal4) | 0.394 | 1.00 |
|  | Adult | (UAS-P3-Sham; *Act5c*-Gal4) to (UAS-P3-RNAi) | < 0.0001 | < 0.001 |
|  |  | (UAS-P3-Sham; *Act5c*-Gal4) to (*Act5c*-Gal4) | < 0.0001 | < 0.0001 |
|  |  | (UAS-P3-RNAi) to (*Act5c*-Gal4) | 0.032 | 0.283 |
| Survival – Fisher’s exact | Pupa | (UAS-P3-RNAi; *Act5c*-Gal4) to (UAS-P3-Sham; *Act5c*-Gal4) | 1.00 | 1.00 |
|  |  | (UAS-P3-RNAi; *Act5c*-Gal4) to (UAS-P3-RNAi) | 0.681 | 1.00 |
|  |  | (UAS-P3-RNAi; *Act5c*-Gal4) to (*Act5c*-Gal4) | 1.00 | 1.00 |
|  |  | (UAS-P3-Sham; *Act5c*-Gal4) to (UAS-P3-RNAi) | 0.692 | 1.00 |
|  |  | (UAS-P3-Sham; *Act5c*-Gal4) to (*Act5c*-Gal4) | 1.00 | 1.00 |
|  |  | (UAS-P3-RNAi) to (*Act5c*-Gal4) | 0.724 | 1.00 |
|  | Adult | (UAS-P3-RNAi; *Act5c*-Gal4) to (UAS-P3-Sham; *Act5c*-Gal4) | < 0.0001 | < 0.0001 |
|  |  | (UAS-P3-RNAi; *Act5c*-Gal4) to (UAS-P3-RNAi) | < 0.0001 | < 0.0001 |
|  |  | (UAS-P3-RNAi; *Act5c*-Gal4) to (*Act5c*-Gal4) | < 0.0001 | < 0.0001 |
|  |  | (UAS-P3-Sham; *Act5c*-Gal4) to (UAS-P3-RNAi) | 0.004 | 0.036 |
|  |  | (UAS-P3-Sham; *Act5c*-Gal4) to (*Act5c*-Gal4) | 0.002 | 0.025 |
|  |  | (UAS-P3-RNAi) to (*Act5c*-Gal4) | 0.868 | 1.000 |

**Table S3.** Raw survival data from larval to pupal (pupa) and pupal to adult (adult) stages. Demonstrates the # or alive or dead, # dead in previous stage, proportion alive, proportion alive from previous stage and # of individuals in that stage.

| Genotype | N | Alive | Dead | Stage | Proportion alive | Proportion of previous |
|---|---|---|---|---|---|---|
| UAS-P3-RNAi; *Act5c*-Gal4 | 100 | 92 | 8 | pupa | 0.92 | 0.92 |
| UAS-P3-RNAi | 216 | 194 | 22 | pupa | 0.8981 | 0.8981 |
| UAS-P3-Sham; *Act5c*-Gal4 | 106 | 97 | 9 | pupa | 0.9151 | 0.9151 |
| *Act5c*-Gal4 | 162 | 148 | 14 | pupa | 0.9136 | 0.9138 |
| UAS-P3-RNAi; *Act5c*-Gal4 | 92 | 0 | 92 | adult | 0 | 0 |
| UAS-P3-RNAi | 194 | 171 | 23 | adult | 0.7917 | 0.8814 |
| UAS-P3-Sham; *Act5c*-Gal4 | 97 | 95 | 2 | adult | 0.8962 | 0.9794 |
| *Act5c*-Gal4 | 148 | 129 | 19 | adult | 0.7963 | 0.8716 |

**Table S4.** Phenotypes displayed at time of recorded death of P3 knockdown group. Phenotypes range from brown puparium/prepupa (1-12h after pupation initiation), developing structures (no pigment; 12-71h), developing pigment (71-90h), developed within puparium (>90h). Note that no P3 knockdown flies eclosed as adults.

| **Stage** | **Counts** | **% of Total** |
|---|---|---|
| brown puparium/prepupa | 29 | 33.33% |
| developing structures | 38 | 43.67% |
| developing pigment | 15 | 17.25% |
| developed | 5 | 5.75% |

## Notes

### Competing Interest Statement

The authors have declared no competing interest.

### Funding

- Natural Sciences and Engineering Research Council

